# Characterizing the Liquid-liquid Phase Co-existence in Biomembrane: Insights from Local Non-affine Deformation and Topological Rearrangements

**DOI:** 10.1101/231274

**Authors:** Sahithya S. Iyer, Madhusmita Tripathy, Anand Srivastava

**Author notes:** Equal contribution (Sahithya S. Iyer, Madhusmita Tripathy).

## Abstract

Lateral heterogeneities in bio-membranes play a crucial role in various physiological functions of the cell. Such heterogeneities lead to demixing of lipid constituents and formation of distinct liquid domains in the membrane. We study lateral heterogeneities in terms of the topological rearrangements of lipids, to identify liquid-liquid phase co-existence in model membranes. By quantifying the degree of non-affineness associated with individual lipid, we are able to characterize the liquid ordered (*L_o_*) and liquid disordered (*L_d_*) phases in model lipid bilayers, without any prior knowledge on chemical identity of the lipids. We explore the usage of this method on all atom and coarse-grained lipid bilayer trajectories. This method is helpful in defining the instantaneous *L_o_-L_d_* domain boundaries in complex multi-component bilayer systems. The characterization can also highlight the effect of line-active molecules on the phase boundaries and domain mixing. Overall, we propose a framework to explore the molecular origin of spatial and dynamical heterogeneity in bio-membranes systems, which can not only be exploited in computer simulation, but also in experiments.

## 1 Introduction

Plasma membrane is a complex self-assembly of a variety of lipids, sterols and proteins. Differential molecular interactions among these diverse constituents gives rise to dynamic lateral heterogeneities in the membrane structure [1, 2]. Such heterogeneities lead to phase separation of the membrane constituents, forming distinctive domains that can be differentiated in terms of their characteristic length and time scales. These domains play vital role in physiological functions of the cell, such as acting as platforms for cell signalling [3], membrane trafficking [4], assembly of protein complexes [5], pathogen uptake [6, 7, 8], and vesicle trafficking [9]. Domain formation in plasma membrane has also been coupled to conformational changes such as domain budding [10]. Most of these cellular functions have been attributed to a special kind of lipid domains known as “rafts”, which are usually perceived as dynamic nano-scale domains, rich in sterol and sphingolipids [11, 1, 12, 2]. However, their existence and size is still debated [13, 14, 15, 16, 17].

Based on the experimental phase diagram of dipalmitoyl phosphatidylcholine (DPPC) and Cholesterol mixture, proposed by Vist and Davis [?], J. H. Ipsen et al. were the first to theoretically characterize the lateral heterogeneities in this multicomponent lipid system [18]. They proposed the coexistence of two de-mixed liquid phases, categorized as “liquid ordered” (*L_o_*) and “liquid disordered” (*L_d_*). Later, based on a theoretical framework, M. Nielsen et al. ascribed the molecular origins of this de-mixing to the “decoupling nature” of cholesterol [19]. The L_o_ phase is characterized by high conformational order (similar to the gel phase (*L_β_*)) and translational disorder (similar to nematic phase in liquid crystals (*L_α_*)), usually comprising of saturated lipids. Absence of the long range intra-molecular conformational order is indicative of the *L_d_* phase. Both these phases have been shown to possess similar translational diffusion of the order of 3*μm*^2^/*s* [20].

Beyond multi-component model lipid systems [21, 22], *L_o_* and *L_d_* phases have also been observed in naturally occurring membranes [23, 24, 25]. Theoretical studies have tried to analyze the physical aspects of such phase separation towards understanding the underlying thermodynamics [26, 27, 28]. These lateral segregations are generally observed in lipid systems containing two different lipid types with well separated melting temperatures *T_m_*, and cholesterol. Lipids with low *T_m_* comprises of unsaturated lipids with a floppy conformation, and thus are found predominantly in the *L_d_* phase. The lateral heterogeneities can be broadly classified into nano- and micro-domains, having dimensions ranging from 10-200 *nm* to a few microns, in model systems. A complete phase separation of the *L_o_* and *L_d_* phases corresponds to the thermodynamic equilibrium of the lipid system. However, the complex non-equilibrium processes occurring in the *in-vivo* cell membrane result in the stabilization of nano-domains over the thermodynamically more favourable micro-domains. Specific lipid composition of the cell membrane close to the critical points [29], role of linactant species [30, 31], intrinsic curvature of constituent lipids generating elastic energy [32], dynamical coupling to the actomyosin cytoskeleton [33, 34, 35, 36], and conformational changes of membrane channel proteins [37] are a few example of such processes. In this work we study the effect of linactants in the stabilization of nano-domains in model systems, which represent “rafts” in real plasma membranes.

Linactants are hybrid lipids with one fully saturated and another unsaturated tail. The resulting molecular structure enables them to increase their compatibility with the two separate phases. These molecules stabilize the nano-domains by reducing line tension in structures (domains) that have a thermodynamically unfavourable high perimeter to area ratio. Line tension is the 2-dimensional analogue of surface tension, present at the interface of two phases, which arises due to the “hydrophobic mismatch” in the thickness of the *L_o_* and *L_d_* phases and membrane curvature [38, 39]. It is defined as the excess energy per unit length of a contact line where two or more distinct phases coexists. Owing to their “line active” nature, linactant molecules preferentially segregate at the interface of the two phases [40], thus lowering the unfavourable interaction energy and thereby, line tension. Line tension is one of the most important determinants of the existence, size, and segregation dynamics of domains in membrane systems [41, 42, 43]. They are known to favour small fluctuating domains and increase their lifetimes [44]. In the present study we have used PAPC as a model linactant molecule, with one tail similar to that of DPPC lipid and the other to that of DAPC lipid.

Our current study is focused on understanding the molecular origin of functionally important phase separation in model lipid systems. Toward this, we employ a characterization method which, while relatively new to biological literature, has been extensively used in the field of amorphous and glassy systems. Though seemingly distinct, amorphous materials have been shown to share striking similarities with biomolecular systems. With functional proteins, they share some fundamental characteristics, in terms of their interior packing [45], rugged free energy surface [46], and relaxation mechanisms [47]. Similar to lipid systems, super cooled glasses have also been found to show dynamic heterogeneities [48, 49]. Motivated by such essential similarities between biological systems and amorphous granular materials, we borrow the idea of non-affine strain measurements, which was used by Falk and Langer in their seminal work [50] to identify the locations of irreversible plastic rearrangements in order to understand the phenomena of viscoplasticity in amorphous solids. In our work, we use this prescription to numerically calculate the degree of non-affine deformation in topological rearrangement of lipids, in their local neighbourhood, to distinguish between *L_o_* and *L_d_* phases. It should be noted that affine and non-affine deformation have previously been employed in granular material [51, 52, 53] and biological systems, in the area of cell mechanics to model actin cytoskeletal networks [54, 55, 56], but in a very different context. To the best of our knowledge, this is the first time that the degree of non-affineness has been used in the context of topological rearrangements to distinguish between the coexisting liquid phases in lipid systems.

Our main results are the following. We show that the degree of non-affineness, of the coexisting phases, is not biased by the chemical identity of the lipid species and serves as a useful tool to distinguish between *L_o_* and *L_d_* phases, and identify the corresponding phase boundary. Using distance based clustering analysis, we show that cholesterol does not preferentially segregate with one of the lipid phase, and rather stations itself at the interface of the two phases. Finally, we also show that the effect of linactant on the liquid phase coexistence can be captured by the spread in the degree of non-affineness of the two separate phases.

The rest of the paper is organized as follows. In Section 2, we discuss our main results. Section 3 details the simulation and characterization methodologies. We conclude in Section 4 with a short summary and possible future directions.

## 2 Results and Discussion

In this section, we present the important results from current study. We begin with an exploration of the proposed numerical method in the context of the existing characterization techniques of *L_o_* and *L_d_* phases in the simulation literature, and then discuss its advantages. Next, the partitioning of cholesterol in such a phase separated lipid systems is investigated in terms of their clustering with the lipid domains. Finally, the effect of linactant molecules on phase boundaries is analyzed in terms of the spread in the degree of non-affineness of the two sub-phases.

### 2.1 Use of non-affine strain measurements to identify *L_o_* and *L_d_* phases

There exists several experimental techniques to identify the two co-existing liquid phases in a lipid system. The most widely used methods involve the measurement of difference in bilayer thickness using AFM [57, 20], and SANS [43], conformational order of the alkyl chain using NMR tail order parameter [58, 59], and diffusion coefficients using FRET [60] and NMR [61]. These methods, along with the calculation of pair correlation functions, have also been traditionally used to study the *L_o_* and *L_d_* phases in simulation literature. The significance of our proposed method, involving non-affine strain measurements, vis-a-vis the merits and pitfalls of the existing methods is discussed in Supporting Information (SI) (Figure S1, S2 and S3).

In this work, we characterize the ordered/disordered regions in the lipid systems in terms of distinction in their topological rearrangements. The details of the formalism is discussed in Materials and Methods section. We quantify this difference by means of the degree of non-affineness, which we define as a measure of the local non-affine strain, associated with individual lipids within a neighbourhood of radius Ω. Taking a particular simulation snapshot as reference, we calculate the residual non-affineness (*χ*^2^) for each lipid in the system at a later time of 10-100 ns. The phosphate head-group of the lipids denote the position of lipids for the calculations and the neighbourhood radius Ω is taken to be 14 Â. The details of the All Atom (AA) and Coarse-Grained (CG) systems, on which the method was tested, are given in Table 1 and Table 2 respectively. As shown in Figure S4 of the SI, the overall value of *χ*^2^ increases monotonically as the time window (Δt) between the two snapshots increases. The average values of *χ*^2^, calculated over the equilibrium trajectories for DAPC/DPPC/CHOL and DUPC/DPPC/CHOL CG systems using a time window of 100 ns and 10 ns of CG time respectively, are shown in Figure 1a. As evident from the figure, *χ*^2^ values show a higher dispersion for the latter systems. This can be attributed to the smaller sizes of domains in DUPC/DPPC/CHOL system, which are formed because of the lower degree of unsaturation of alkyl chain of DUPC, as compared to DAPC. DAPC/DPPC/CHOL forms continuous domains which is reflected in the lesser dispersion of *χ*^2^ values. The spread in *χ*^2^ values can also give a qualitative information about the correlation decay length *ξ*, which is a measure of the size of fluctuations of the lipid domains [62, 63]. Higher values of *ξ*, indicative of larger wavelength fluctuations [64], can be associated with a high dispersion in *χ*^2^ values. It should be emphasized that the calculation of *χ*^2^ values is computationally less expensive in comparison to the estimation of *ξ* from the decay of spatial correlation function. Figure 1b shows the spread in *χ*^2^ values for single phase *L_o_* and *L_d_*, and mixed phase *L_o_*/*L_d_* systems from tens of microseconds long AA trajectories. Systems (A) and (B) have a unimodal distribution in *χ*^2^ values. This indicates the presence of a single phase. The magnitude of *χ*^2^ values can be used to distinguish between *L_o_* and *L_d_* phases. System (C) is phase separated and shows a bimodal distribution in *χ*^2^ values. PSM lipids with higher *χ*^2^ values can be mapped back to those lipids which are not part of the continuous PSM domain, and are interspersed with the DOPC lipids (image not shown). Hence, this method identifies *L_o_* and *L_d_* phases without being biased by the chemical identity of the lipids.

**Figure 1a:**
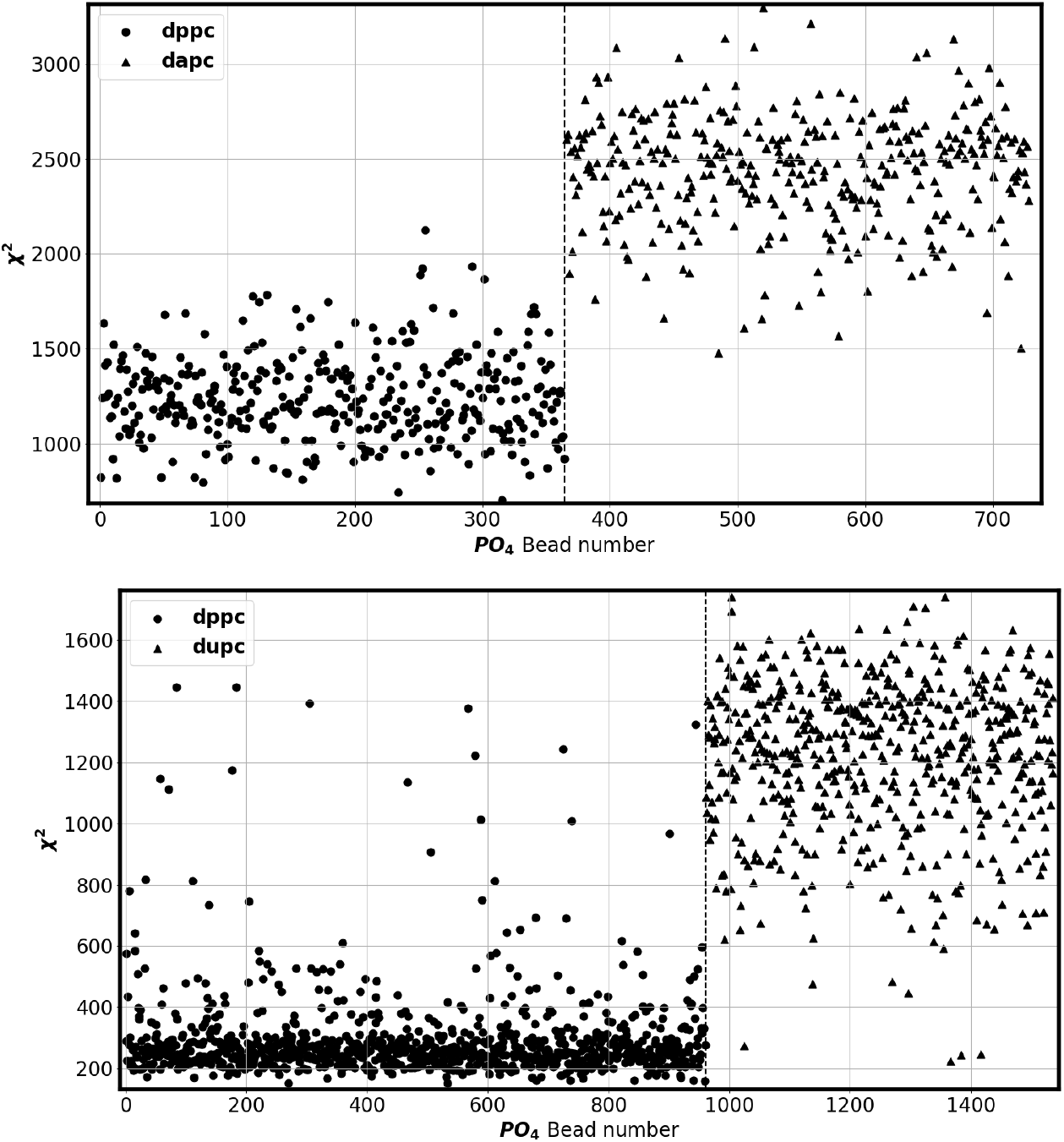
Bimodal distribution of *χ*^2^ values calculated for the *PO*_4_ groups for (top) DAPC/DPPC/CHOL system and (bottom) DUPC/DPPC/CHOL system. Figure (S4) in SI shows that the difference in *χ*^2^ values for these two phases increases linearly with time both for CG and AA systems.

**Figure 1b:**
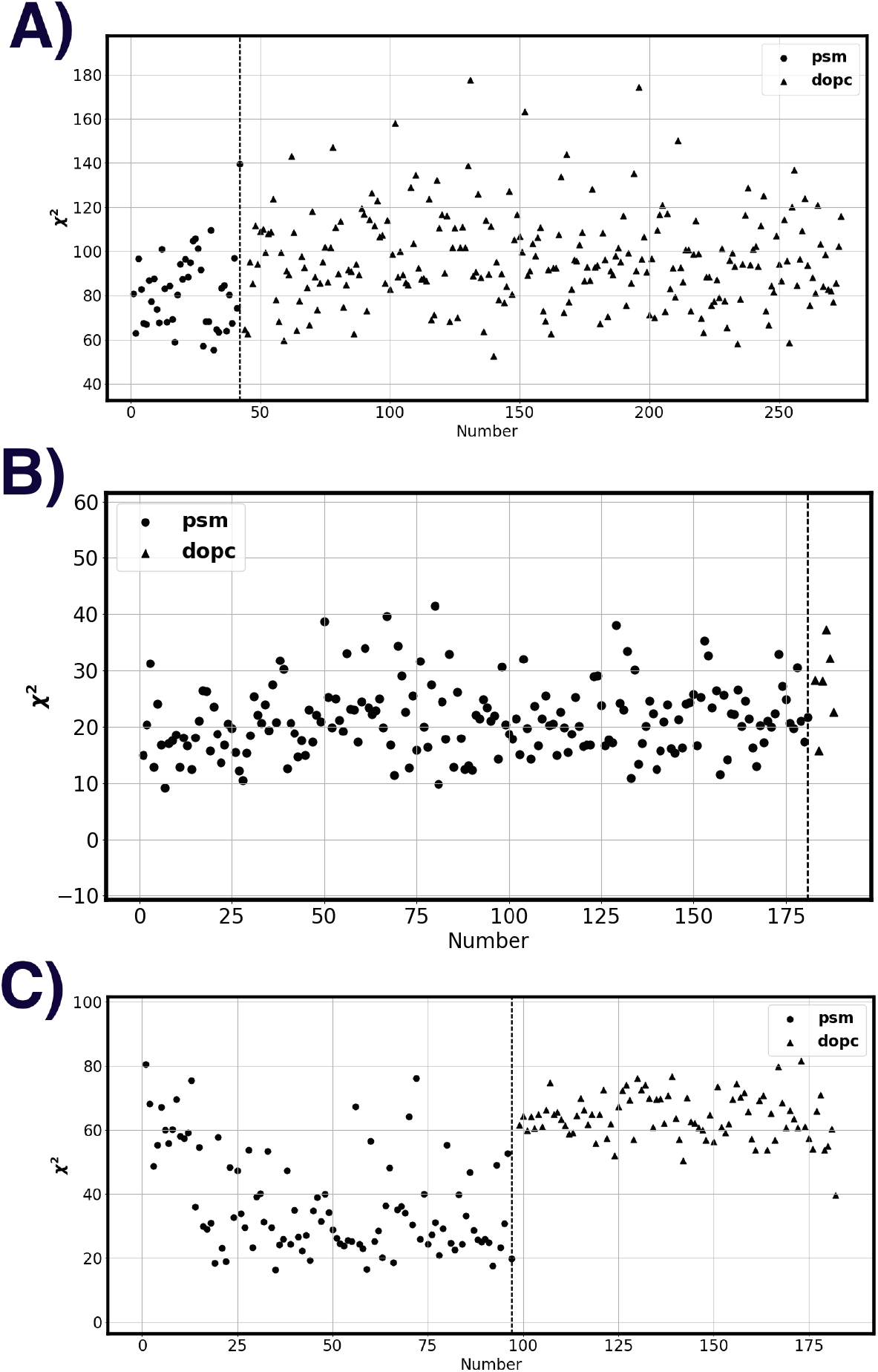
Plot of *χ*^2^ values calculated for the P atoms for various systems. (A) PSM/DOPC/CHOL composition control *L_d_* system (B) PSM/DOPC/CHOL control *L_o_* system having uniformly distributed *χ*^2^ values and (C) PSM/DOPC/CHOL mixed *L_o_/L_d_* system captured by the bimodal distribution in *χ*^2^ values.

**Table 1:** Simulation details of various all atom trajectories analyzed in this work.

| All Atom Systems |  |  |  |  |  |
| --- | --- | --- | --- | --- | --- |
| Lipid system | Composition | Phase | Temperature (K) | System size (nm) | Duration ($\mu s$) |
| PSM/DOPC/CHOL | 0.15/0.82/0.03 | Ld | 295 | $13.42 \times 13.42$ | 0.81 |
| PSM/DOPC/CHOL | 0.64/0.03/0.33 | Lo | 295 | $10.63 \times 10.63$ | 0.95 |
| PSM/DOPC/CHOL | 0.43/0.38/0.19 | Lo/Ld | 295 | $10.86 \times 10.86$ | 4.54 |
| PSM/POPC/CHOL | 0.61/0.08/0.31 | Lo | 295 | $10.62 \times 10.62$ | 9.08 |
| PSM/POPC/CHOL | 0.47/0.32/0.21 | Lo/Ld | 295 | $10.87 \times 10.87$ | 6.88 |
| DPPC/DOPC/CHOL | 0.37/0.36/0.27 | Lo/Ld | 298 | $9.53 \times 9.53$ | 9.43 |

**Table 2:** Simulation details of the CG systems (Martini).

| Coarse grain systems |  |  |  |  |
| --- | --- | --- | --- | --- |
| Lipid system | Composition | Phase | Temperature(K) | System size(nm) |
| DAPC/DPPC/CHOL | 0.35/0.35/0.30 | Lo/Ld | 295 | 24.82×24.82 |
| DUPC/DPPC/CHOL | 0.25/0.42/0.33 | Lo/Ld | 290 | 35.5×35.5 |

Recently, Hidden Markov Model (HMM) was used to identify *L_o_* and *L_d_* phases in AA simulation of DPPC/DOPC/CHOL system [65]. To benchmark our analysis, we identify the liquid phases in the same AA systems using *χ*^2^ values. To be consistent with earlier results, the mid-carbon sites of each of the lipid tails are used as evolving coordinates for the *χ*^2^ analysis. Figure 2a shows a color map of the lipid coordinates in the AA system, with *χ*^2^ values as the colorbar. The ordered *L_o_* nano-domain can be identified as a patch with the lowest *χ*^2^ values and is mostly composed of DPPC lipids, in agreement with the previous HMM analysis (Movie S1 in SI). The disordered *L_d_* phase is associated with comparatively higher *χ*^2^ values. It should be stressed here that the *χ*^2^ analysis does not require the information about the chemical identity of the lipids/cholesterol and is purely based on the topological data. As with PSM lipids with higher *χ*^2^ values (Figure 1b C), there are also DPPC lipids with higher *χ*^2^ values and DOPC lipids with lower *χ*^2^ values. Overall, this formulation allows us to capture the ordered/disordered micro-domains in CG and nano-domains in AA simulation trajectories.

**Figure 2:**
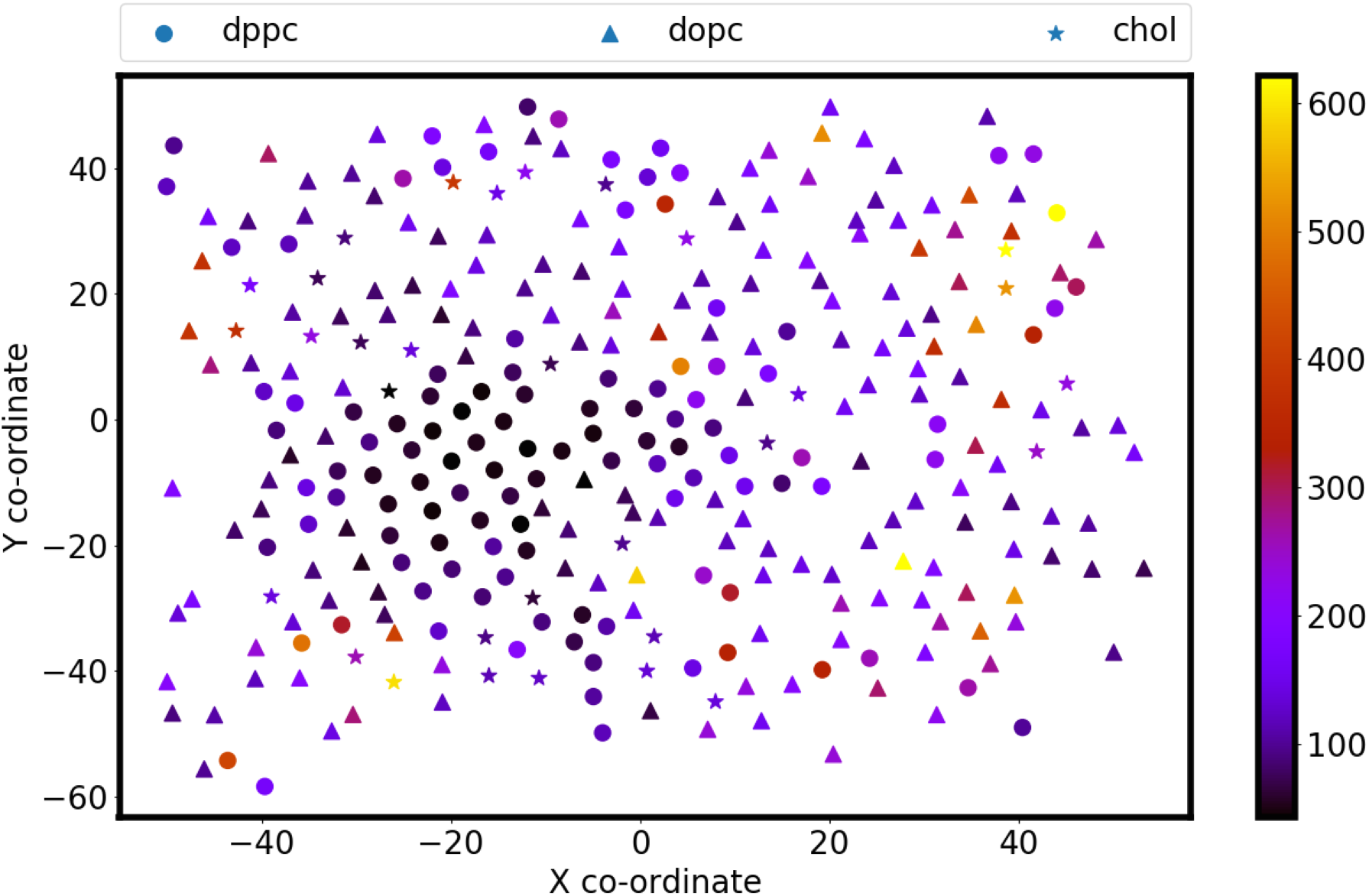
Position of DOPC, DPPC and CHOL mid atom sites color coded with the corresponding *χ*^2^ values is show. DPPC lipid sites shown as circles for a nanodomain which is captured by the low *χ*^2^ values. Other DPPC lipid sites that are not part of this continuous domain have higher *chi*^2^ values.

Beyond the ability to discern between ordered/disordered domains, *χ*^2^ values can also be used to monitor the kinetics of domain formation within the limitation of the CG trajectories. The initial stage of phase separation has been proposed to occur though spinodal decomposition near the critical point and via nucleation otherwise [66]. Figure 3 shows the long tail distribution of *χ*^2^ values for DAPC and DPPC lipids in CG DAPC/DPPC/CHOL system as the simulation advances. For this part of the calculations, the values of *χ*^2^ have been segregated based on the lipid type and the distributions are calculated separately. The overlapping region in the distribution corresponds to the lipids that do not belong to a pure *L_o_* or *L_d_* phase. As apparent from the figure, the overlap in the distribution decreases with time indicating separation of the two phases, and thus can be used as a measure to study the segregation dynamics. As shown in Figure 4, as the area of the overlapping region converges the system is said to have attained equilibrium.

**Figure 3:**
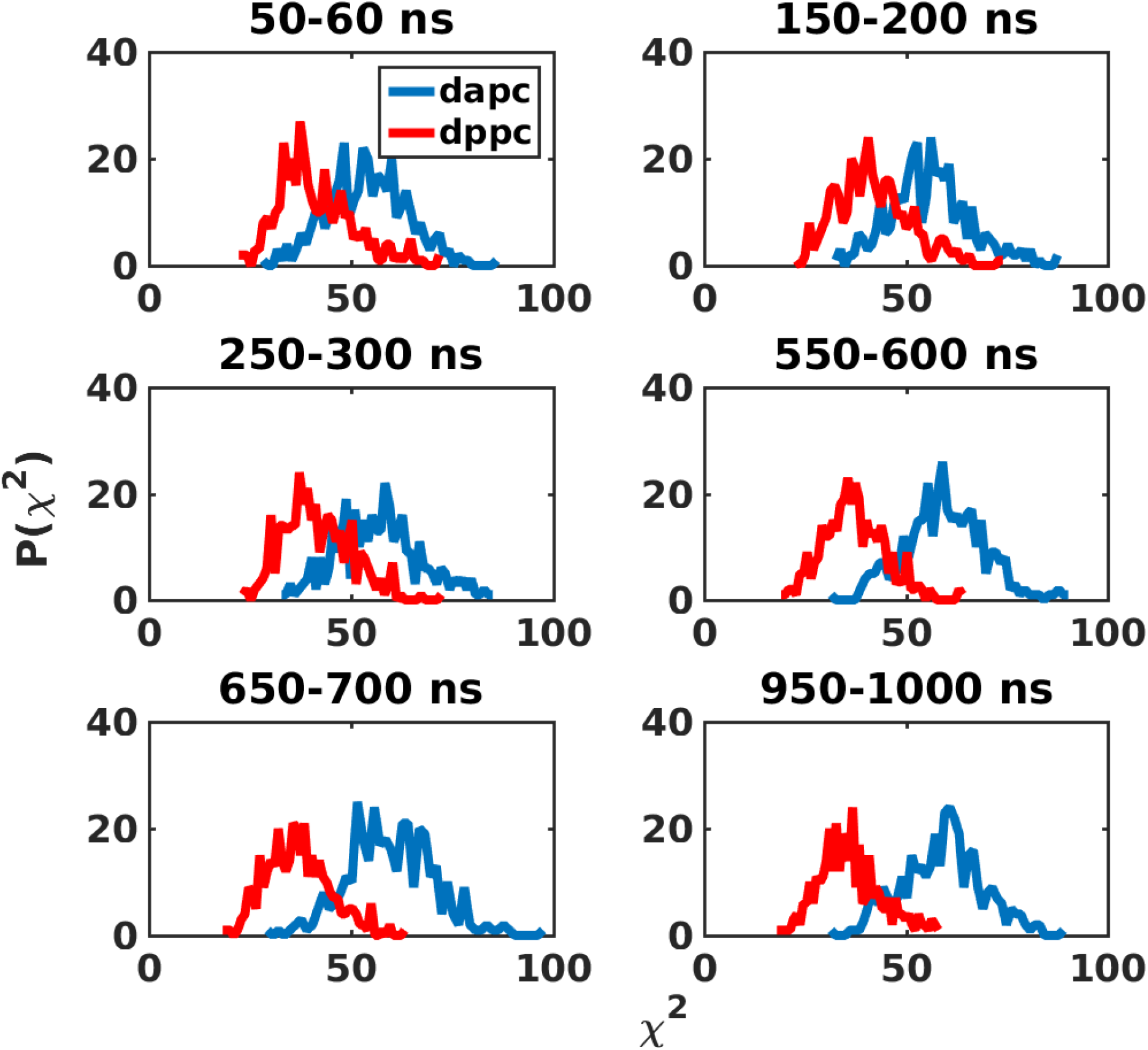
*χ*^2^ values capturing dynamics of phase segregation in DAPC/DPPC/CHOL system.

**Figure 4:**
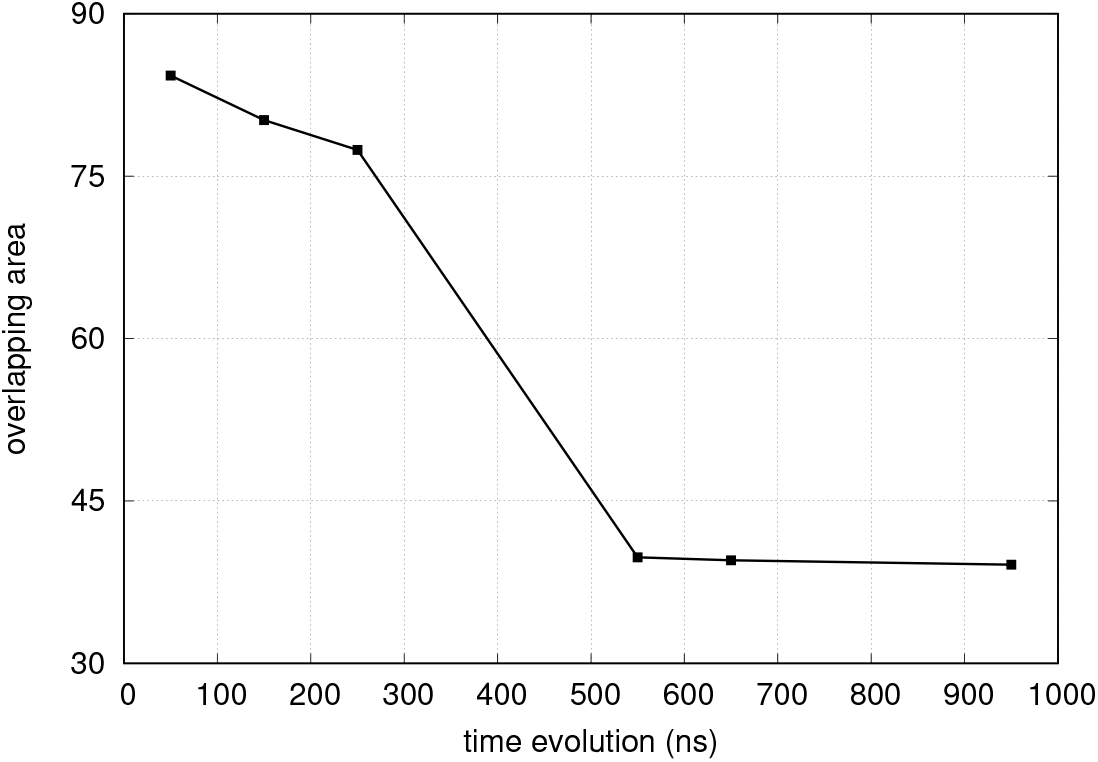
The evolution of area of the overlapping region of the long tails of *χ*^2^ distributions can be used to monitor domain growth.

### 2.2 Identifying the “true” interface of ordered/disordered regions

In general, traditional way of identification of the boundary lipids involves the association of saturated and unsaturated lipids to different phases [18]. The color map in Figure 2 for AA DOPC/DPPC/CHOL system shows that not all DPPC lipids form *L_o_* phase as previously assumed in HMM calculations [65], rather some DOPC and cholesterol sites also belong to *L_o_* phase. Our calculations thus show that “local order” is environment dependent and need not be associated with the chemical identity of the lipids. As seen in Figure 2a, some of the DPPC lipids have higher *χ*^2^ and these can be rightly identified as those belonging to the *L_d_* phase. The *χ*^2^ values are shown in Figure 2b for clarity. Lower S_CD_ of these DPPC lipids with higher *χ*^2^ values also shows that they are less ordered. In Figure 5, shown are the contoured heat maps of the DAPC/DPPC/CHOL and DUPC/DPPC/CHOL CG systems, with averaged *χ*^2^ values as the colorbar. As evident, the boundary lipids acquire intermediate *χ*^2^ values, and thus can be used to detect lipids that form the interface of *L_o_* and *L_d_* phases. The sites with intermediate *χ*^2^ values are shown for the DUPC/DPPC/CHOL CG system in Figure S5. These sites trace out the *L_o_* domain boundary of the system. Further the fluctuation in this domain boundary can be used to quantify line tension in the system.

**Figure 5:**
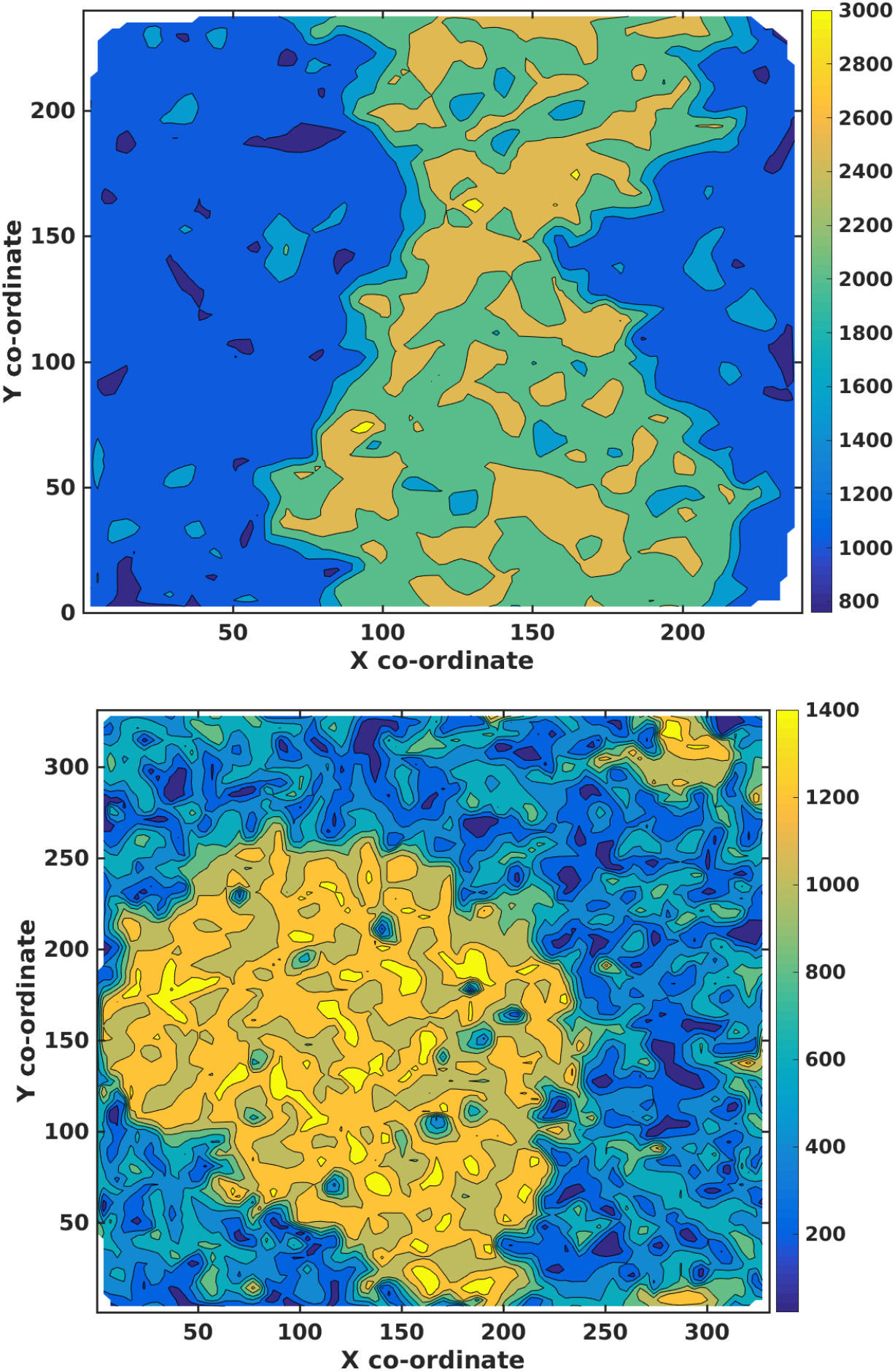
Contour map with *χ*^2^ has the contour height shows for (top) DAPC/DPPC/CHOL system that lipids with *χ*^2^ values between 45-55 and (bottom) for DUPC/DPPC/CHOL system lipids with *χ*^2^ values between 700-900 can be identified as the boundary lipids.

### 2.3 Cholesterol partitioning in *L_o_* and *L_d_* phases

As discussed in the last section, the ordered phase in the membrane is not formed exclusively by the saturated lipids, but is also composed of some unsaturated lipids and cholesterol. Therefore, to study the behaviour of cholesterol in the phase separated system of ordered and disordered lipids, we perform cluster analysis on the atomistic trajectories of DOPC/DPPC/CHOL, DOPC/PSM/CHOL and POPC/PSM/CHOL systems. The mid points of the two tails of each lipid are taken as the evolving coordinates. A distanced based clustering algorithm is used with cutoffs 5, 6, 7, and 8 Å. The variation in N_S_, the occurrence probability (number of clusters with size S), with different cutoff distances for DPPC (/PSM) and DOPC (/POPC) lipids, is shown in Figure 6, Figure S6 and Figure S7. For both the lipid types, we also calculate the cluster size distribution by taking the cholesterol sites into account. For a given cutoff distance for both DOPC and DPPC lipids, N_S_ increases when the cholesterols are included in the analysis. The occurrence probability of larger clusters (S > 100) increases noticeably for the two lipid types. Such a common trend of increase in N_S_ for both the lipids upon incorporating the cholesterol sites in the calculation, especially for large S, indicates the formation of large clusters of DOPC and DPPC lipids that also include cholesterols. This observation suggests that cholesterol does not segregate preferentially with either of the lipid types, but rather is located at the interface of the DPPC/DOPC clusters. This is in line with the previous study [65], which proposed that the cholesterol does not disrupt the hexagonal packing of DPPC lipids and hence are stationed at the interface of DPPC and DOPC lipid domains.

**Figure 6:**
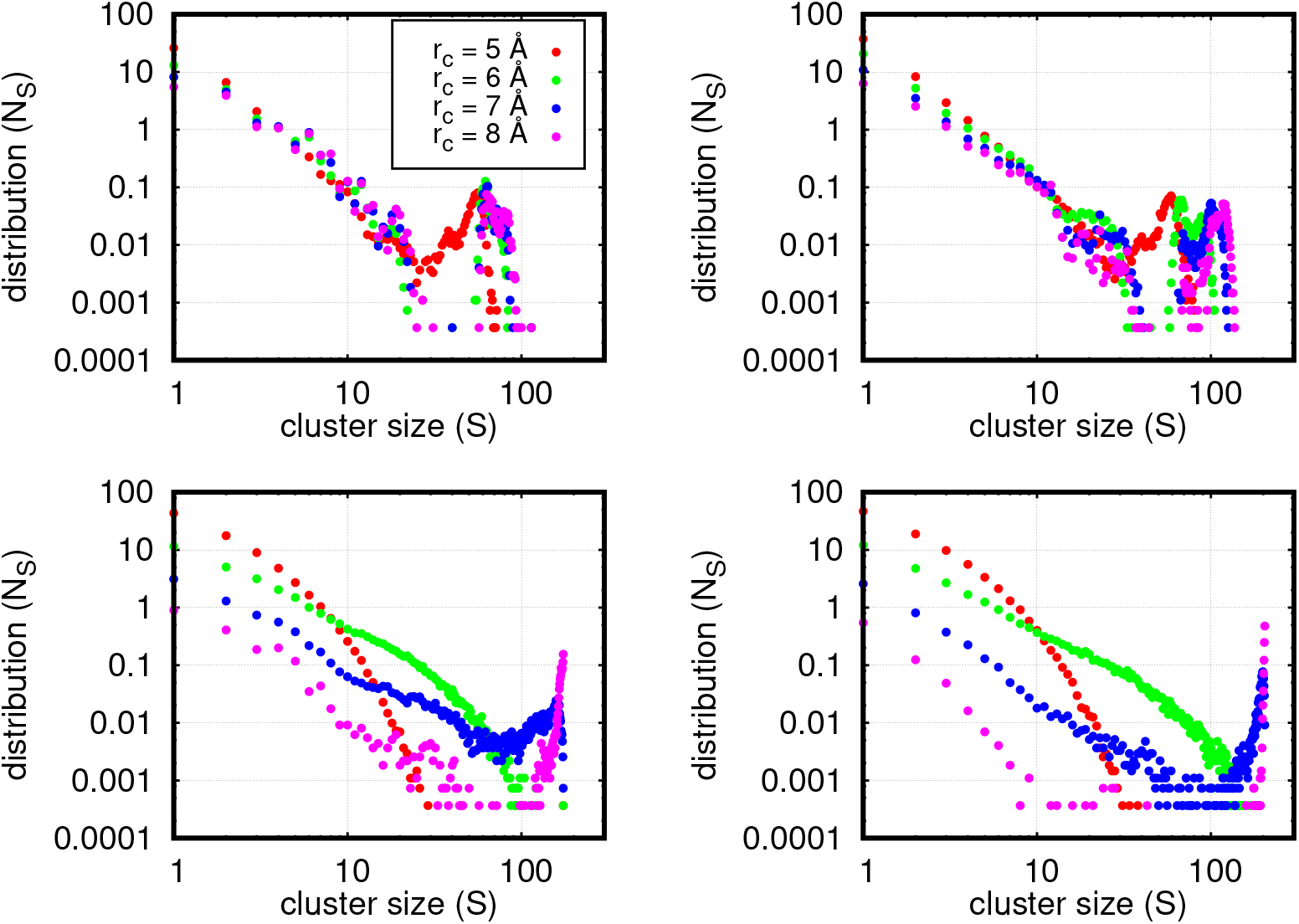
Cluster size distribution for (Top) DPPC and (Bottom) DOPC lipids without (left) and with (right) CHOL, using various cut-off distances.

### 2.4 Predicting the effect of linactant molecules on phase boundaries using *χ*^2^ values

Theoretical study shows that line tension depends quadratically on thickness difference between the *L_o_* and *L_d_* phases [67]. Linactant molecules lower the line tension across the boundary of the two phases by reducing the hydrophobic mismatch [68]. To study the effect of linactant molecules on the phase segregation in DAPC/DPPC/CHOL system, we compute the distribution of *χ*^2^ values for both the lipid types with increasing linactant (PAPC) ratio (X: number of PAPC lipids to total number of unsaturated lipids), as shown in Figure 7. As discussed in earlier sections, the overlap in the long tail probability distribution indicates the lipids at the interface of *L_o_* and *L_d_* phases, and thus can be monitored as an indication of disruption of the phase boundary and increased thermal fluctuations. We find that the overlap in *χ*^2^ distribution monotonically increases with increasing linactant ratio, suggesting increase in the domain boundary fluctuations with increasing number of “interfacial” lipids. Eventually, beyond a certain concentration of these line active molecules, the phase segregation is almost lost as the saturated and unsaturated lipids start to mix without an energy penalty.

**Figure 7:**
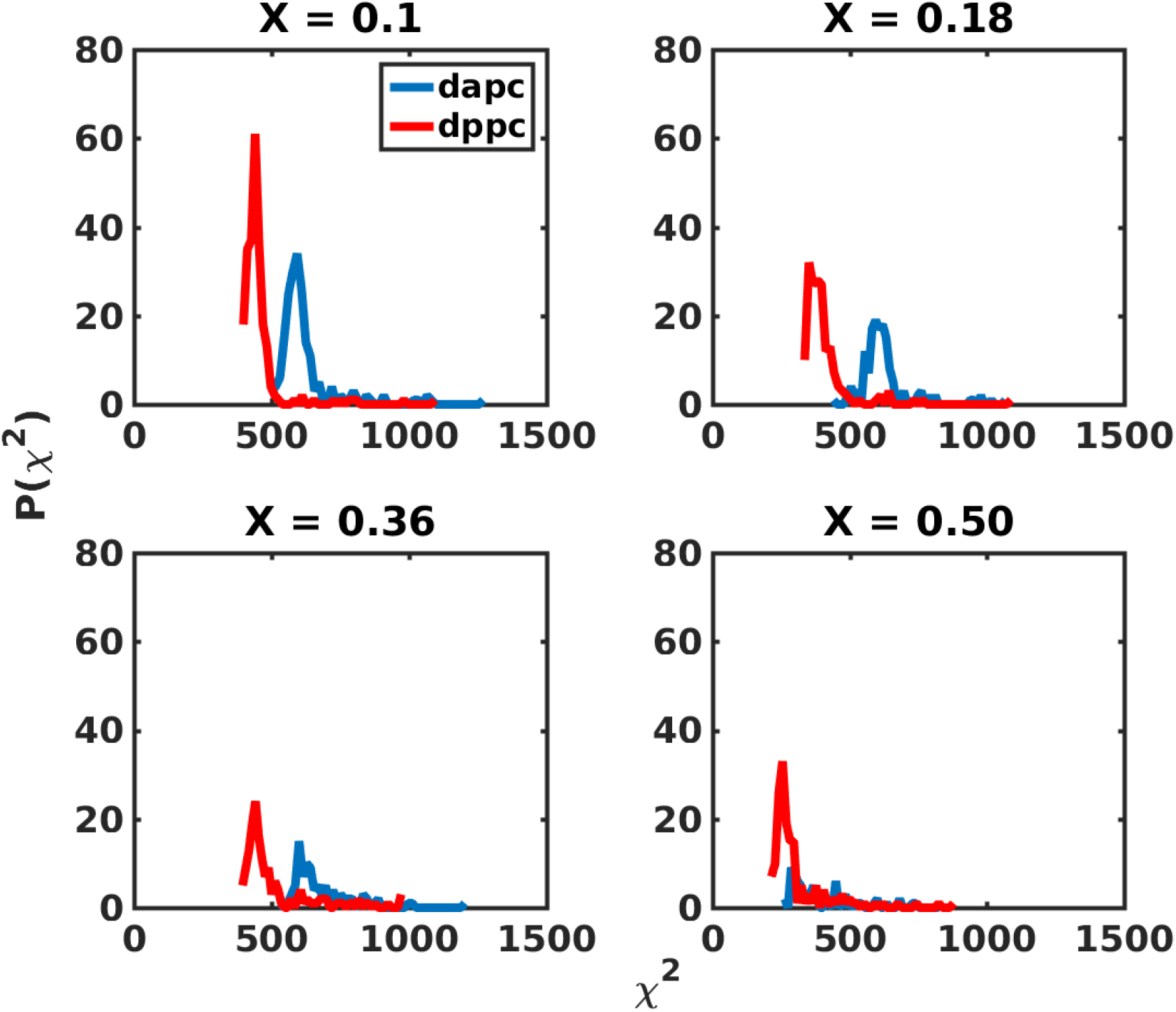
Probability distribution of *χ*^2^ values of DAPC and DPPC lipid types for systems with increasing linactant concentrations shows that the overlap in the long tails increases linearly.

Figure 7 also indicates the *L_d_* phase to be more affected by the linactant molecules, than the *L_o_* phase. The dispersion of *χ*^2^ values, in the *L_d_* phase, increases with increase in linactant ratio, as indicated by the lowering of the peak and its subsequent flattening. This inference is similar to the analysis performed by Rosetti et al. [68] on the same lipid system. They show that the affinity of the saturated tail of PAPC towards the *L_o_* region decreases with increase in PAPC ratio, thus having more effect on the unsaturated DAPC lipids. We calculate the probability distributions of tail order parameter (*S_CC_*) of DAPC and DPPC lipids, for all the linactant ratios probed in their study, which is shown in Figure 8. While the *S_CC_* distributions of DPPC lipids do not change significantly with linactant ratio, those of DAPC lipids change remarkably from a single peak to a wider distribution. The average order parameter values for DPPC changes from 0.611 to 0.554, and that for DAPC from 0.094 to 0.053 for X=0.1 to 0.5. Linactants act by first increasing the disorderedness of DAPC lipids upto a critical concentration of X=0.36 and then ordering them. This trend is also seen in Figure 8. After X=0.36, a finite fraction of the ordered lipids become floppy while some of the disordered lipids become linear, as can be inferred from the spread in the corresponding distributions. Such a redistribution leads to a decrease in the thickness mismatch between the two lipid domains, thereby decreasing the line tension.

**Figure 8:**
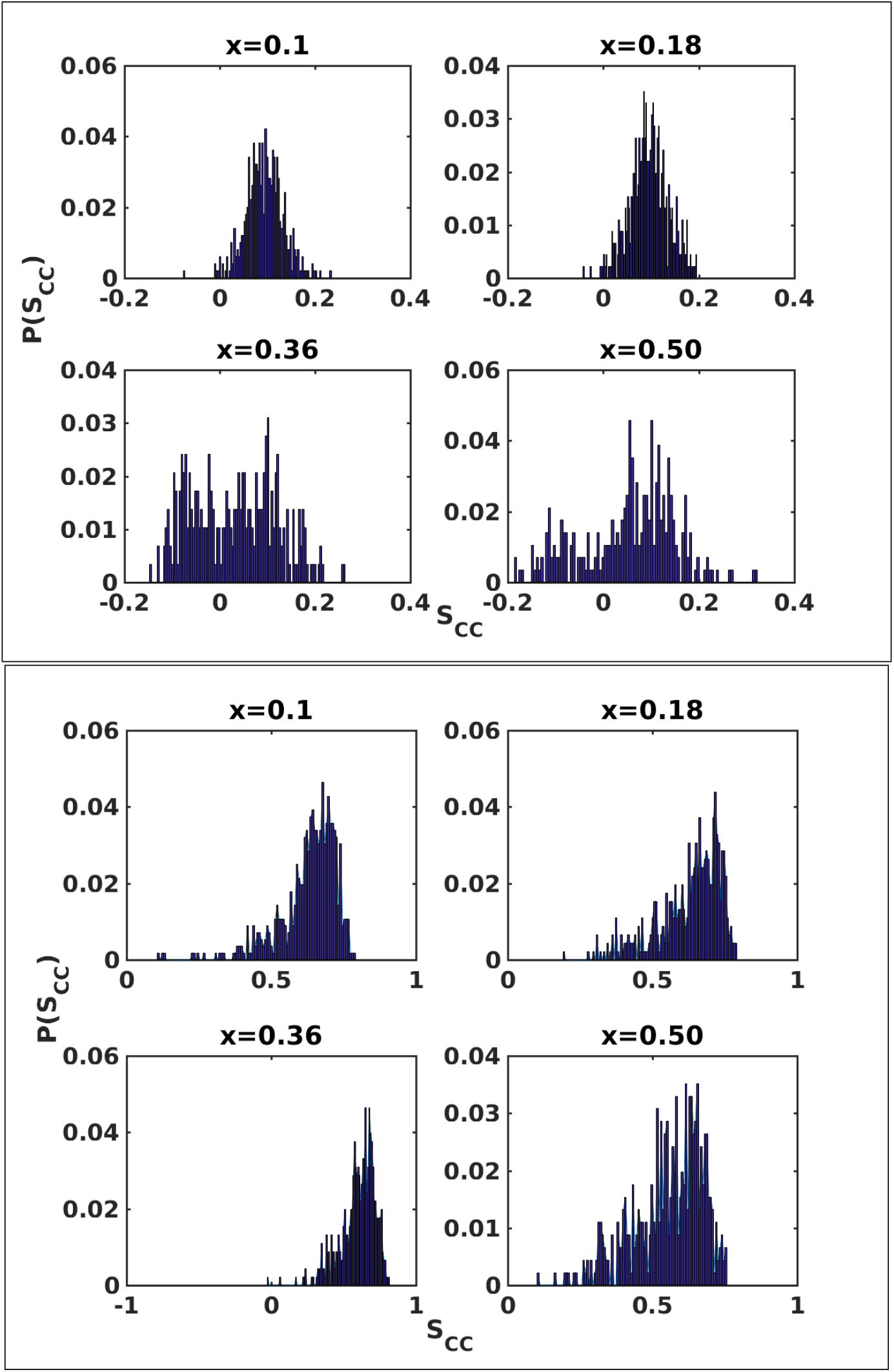
Probability distribution of tail order parameters for (Top) DAPC and (Bottom) DPPC lipids for different linactant ratios.

## 3 Materials and Methods

In this section we summarize the details of the models, simulation methodologies, and numerical implementation. We start by highlighting the details of AA and CG simulations in the first subsection. We then discuss various characterization methods used in this work, including the mathematical construct, we followed, to calculate *χ*^2^.

### 3.1 Model and Simulation details

#### 3.1.1 All Atom Simulations

A diffusion constant of the order of 10^−7^*cm*^2^/*s* for lipids in all atom representation requires the systems to be evolved for several microseconds in order to achieve equilibrium lateral distribution. Therefore, such simulations are computationally expensive. The all atom simulation trajectories used in this work were borrowed from Edward Lyman’s group in Delaware. The trajectories constituted six ternary lipid bilayer simulations performed on Anton supercomputer. Position and velocities were collected at an interval of 240 ps. The relevant details of the all atom systems are summarized in Table 2. Further technical details of the simulation set up can be found in the corresponding references [69, 65].

#### 3.1.2 Coarse Grained (CG) simulations

CG simulation on 35%/35%/30% DAPC/DPPC/CHOL ternary lipid system, of size ~ 25 × 25 Å^2^, was set up using CHARMM-GUI Martini membrane builder web server [70, 71]. The tensionless membrane was simulated using GROMACS 5.0.2 [72] in NPT ensemble at 295 K temperature and 1 bar pressure. A constant temperature was maintained using velocity rescaling and a constant pressure using Parrinelo-Rahman barostat [73] with relaxation times of 1.0 ps and 2.0 ps respectively. Cut-off and shift distances of 1.2 nm and 0.9 nm to 1.2 nm respectively were used for van der Waals interactions. To simulate a tensionless bilayer, semi-isotropic pressure coupling was performed independently in the lateral and normal directions with a compressibility constant of 3 × 10^−4^ bar^−1^. The restraints, initially applied on the lipid head-groups in order to maintain the bilayer structure, were gradually released in the equilibration phase. The multi-component bilayer system was then evolved for 1.6 *μs* and positions and velocities were collected at a frequency of 40 ns, until equilibrium phase separation was achieved.

The CG trajectory for martini coarse-grained DUPC/DPPC/CHOL lipid mixture was obtained from Peter Tieleman’s group at Calgary [74]. The system was evolved for 11 *μs* and positions and velocities were collected at every 10 ns. Trajectories for systems with different linactant fractions were borrowed from Carla M. Rosetti’s lab at Argentina [68]. Positions and velocities of these systems were written at a frequency of 60 ns. Details of all the three CG systems are summarized in Table 3 and 4.

**Table 3:** Simulation details of various CG ternary lipid systems with linactant (Martini).

| Effect of linactant |  |
| --- | --- |
| Hybrid lipid | $x = \frac{Hybrid}{Hybrid+Symmetric}$ |
| PAPC | 0 |
| PAPC | 0.1 |
| PAPC | 0.18 |
| PAPC | 0.36 |
| PAPC | 0.5 |

### 3.2 Numerical Implementation

We calculate the degree of non-affineness, associated with any lipid in an evolving membrane system, based on the topological rearrangements occurring in a local neighborhood around it [50]. An affine deformation is always associated with a uniform strain. For non-affine case, the total deformation can be broken down to an affine and a residual part. Thus, the degree of non-affineness (*χ*^2^) measures the residual non-affine content of a deformation. Following the mathematical construct of Falk and Langer [50], the residual deviation from affine deformation during a time interval [*t*, *t* + Δ*t*] is given by

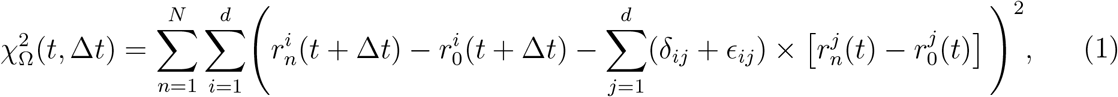

where the indices *i* and *j* run through the spatial coordinates for dimension *d* and *n* runs over the *N* lipids in the neighbourhood Ω, defined within a cutoff distance around the reference lipid *n* = 0. *δ_ij_* is the Kroneker delta function. *ϵ_ij_* is the strain associated with the maximum possible affine part of the deformation and thus, minimizes 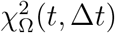 and 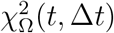 can be calculated easily following the prescription given in the original reference.

For all the calculations the head-group sites were used to as reference to represent a lipid. In order to compare results from AJ Scodt et al. [65], the mid tail sites were used in the case of DOPC/DPPC/CHOL AA system. The calculations on AA systems were performed with Δ*t* = 240 ps. For DUPC/DPPC/CHOL CG systems, Δ*t* = 100 ns in CG timescales and Δ*t* = 4 ns for DAPC/DPPC/CHOL CG systems. For DAPC/DPPC/CHOL systems with varying linactant concentrations Δ*t* = 60 ns was used.

In addition to the degree of non-affineness calculation, analyses traditionally used in the simulation literature on lipid systems to distinguish between *L_o_* and *L_d_* phases, including the calculation of radial distribution functions, tail order parameter, diffusion co-efficient from mean square displacement, thickness and area per lipid, were carried out. The ability of these measurements in characterizing the *L_o_* and *L_d_* phases was investigated and compared to that of non-affineness calculation.

## 4 Summary and Conclusion

In this work, we explore the feasibility of utilizing the information on topological rearrangement of lipids, to characterize the phase separation in a model membrane system. We numerically calculate the degree of non-affineness (*χ*^2^), associated with evolving lipids in their local neighbourhood, to distinguish between the liquid ordered and liquid disordered phases.

Apart from identifying these phases, *χ*^2^ values can also be used to monitor the kinetics of domain segregation. Such a kinetic study can distinguish between the two types of phase separation: nucleation and growth, and spinodal decomposition, depending on the nature of domain evolution. The non-affine parameter contains information about the local neighbourhood of the lipids and hence can be used to identify lipids at the boundary of the two phases. Unlike the existing methods that are traditionally used in simulation studies and experiments, the boundary thus traced out is not biased by chemical identity of the lipids and so identifies the “true interface”. The effect of linactant molecules that preferentially segregate at the interface of *L_o_* and *L_d_* regions can be captured using the overlap between the probability distribution of *χ*^2^ values for lipids in both phases. We show that linactant molecules lower the line tension by decreasing hydrophobic mismatch between the lipids that form the interface of the two phases.

On the methodology front, we propose a computationally less expensive method of characterizing *L_o_* and *L_d_* phases in atomistic and coarse-grained simulation studies of bio-membranes. This method can also be utilized in kinetics experiments, where images separated by timescales as large at 0.5 *μs* can be analyzed without performing complicated image processing. While identification of individual lipids in experiments can be severely limited by the experimental resolution, one can nonetheless utilize this method in the study of kinetics of large-scale membrane features such as lipid rafts and protein assembly, and tissues.

## 5 Acknoledgement

The authors thank Edward Lyman, Peter Tieleman, and Carla M. Rosetti’s lab for sharing the simulation trajectories. The financial support from Indian Institute of Science-Bangalore and the HPC facility “Arjun”, setup from grants by Department of Biotechnology (DBT-India), are greatly acknowledged.

